# Sense-antisense gene overlap causes evolutionary retention of the few introns in *Giardia* genome and the implications

**DOI:** 10.1101/333310

**Authors:** Min Xue, Bing Chen, Qingqing Ye, Jingru Shao, Zhangxia Lyu, Jianfan Wen

## Abstract

**Background:** It is widely accepted that the last eukaryotic common ancestor (LECA) and early eukaryotes were intron-rich and intron loss dominated subsequent evolution, thus the presence of only very few introns in some modern eukaryotes must be the consequence of massive loss. But it is striking that few eukaryotes were found to have completely lost introns. Despite extensive research, the causes of massive intron losses remain elusive, and actually the reverse question – how the few introns are retained under the pressure of loss is equally significant but was rarely studied, except that it was conjectured that the essential functions of some introns prevent their loss. The extremely few (eight) spliceosome-mediated cis-spliced introns in the relatively simple genome of *Giardia lamblia* provide an excellent opportunity to explore this question.

**Results:** Our investigation of the intron-containing genes and introns in *Giardia* found three types of intron distribution patterns: ancient intron in ancient gene, relatively new intron in ancient gene, and relatively new intron in relatively new gene, which can reflect to some extent the dynamic evolution of introns in *Giardia*. Not finding any special features or functional importance of these introns responsible for the retention, we noticed and experimentally verified that some intron-containing genes form sense-antisense gene pairs with functional genes on their complementary strands, and that the introns just reside in the overlapping regions.

**Conclusions:** In *Giardia*’s evolution, despite constant pressure of intron loss, intron gain can still occur in both ancient and newly-evolved genes, but only a few introns have been retained; the evolutionary retention of introns is most likely not due to the functional constraint of the introns themselves but the causes outside of introns, such as the constraints imposed by other genomic functional elements overlapping with the introns. These findings can not only provide some clues to find new genomic functional elements -- in the areas overlapping with introngs, but suggest that “functional constraint” of introns may not be necessarily directly associated with intron loss and gain, or that the real functions or the way of functioning of introns are probably still outside of our current knowledge.

## Background

Spliceosomal introns are a common feature of all eukaryotic nuclear genomes, but their number and density in a genome vary dramatically among different species [1, 2], ranging from less than 0.5 intron/gene in some protists such as the Microsporidian *Encephalitozoon* species [3] and *Cyanidioschyzon merolae* [4] to over 18 per gene in *Symbiodinium minutum* [5] (even larger than those of most mammals, which is generally over eight/gene [6]). Accumulating evidence suggests that the LECA and early eukaryotes were relatively intron rich, with subsequent genome evolution dominated by intron loss, and thus those contemporary eukaryotes with remarkably few introns must have experienced massive intron loss secondarily [7–9]. But, interestingly, no eukaryotes with sequenced genomes so far have been found to have completely lost their introns except the two Microsporidia species, *Nematocida parisii* and *Nematocida* sp1[10].

Unfortunately why introns were lost, especially massively lost in some eukaryotes, remains obscure despite extensive research [11, 12]. Obviously, the reverse question -- how introns, especially the few ones in intron-poor eukaryotes, can be retained under the pressure of loss is equally important, but it was rarely carefully studied. Although some authors thought that the reason for the retention of introns in genomes is likely due to the essential functions of these introns [13–15], this ‘functional constraint’ scenario – “only the introns with important functions can get rid of the fate of being lost” lacks evidence that the lost introns are all useless or less useful than the retained ones in any eukaryotes, and moreover, actually the functions of introns are still far from being well understood [16]. Therefore, the investigation of the evolutionary retention of intron might be helpful not only to answering the question about intron loss but also to understanding the function and evolution of introns.

*Giardia lamblia* is a parasitic protozoan belonging to Diplomonadida (Excavata). It has a very minimalistic genome, compact in structure and content [17], and only eight spliceosomal introns were found in its genome [17–22]. Thus it can be speculated that this organism must have undergone massive intron loss, with very few left in the genome. Therefore, this organism may provide an excellent opportunity for exploring how the few introns were retained. In the present work, by investigating the intron-containing genes and the few introns of *Giardia*, besides finding the distribution patterns that can reflect the dynamic evolution of intron in *Giardia*, we observed and experimentally confirmed an interesting phenomenon that sense-antisense (SAS) gene overlaps appear in the areas of some introns, and thus “overlap constraint” might be at least one of the causes for preventing introns from being lost, though it is uncertain whether the other retained introns also overlap with any unknown genomic functional elements yet. The implications of these findings for intron evolution and function are discussed.

## Results

### Characteristics of the intron-containing genes and their introns in *Giardia* genomes

In the genome database of *Giardia*, GiardiaDB, four of the eight *Giardia* intron-containing genes are annotated to code proteins with sequence similarity to known proteins, and the other four to code hypothetical proteins. Our investigation (mainly by sequence comparative analysis) indicated that: 1) the former four intron-containing genes are all common eukaryotic-conserved genes, which are most likely vertically inherited from the LECA and thus are very ‘ancient’, while the latter four are all Giardia-specific genes (not found in other eukaryotes including other Excavata species), which thus most likely emerged after the divergence of *Giardia* from other Excavata and thus are ‘relatively new’ genes compared with the ‘ancient’ ones above; 2) the introns in the three (GL50803-15604, GL50803-15124, GL50803-17244) of the four ancient genes are eukaryotic-conserved (Additional file 1), and thus they are ‘ancient’ introns in ancient genes, while the intron in the other one ancient gene (GL50803-27266) is a *Giardia*-specific intron (not found in other eukaryotes including other Excavata species), and thus this intron most likely emerged after the divergence of *Giardia* from other Excavata and is a ‘relatively new’ intron in an ancient gene; 3) all four Giardia-specific (‘relatively new’) intron-containing genes (GL50803-37070, GL50803-35332, GL50803-15525 and GL50803-86945), which account for only about 0.6 percent of all the ~700 *Giardia-specific* protein-coding genes in the genome [17], each contain an Giardia-specific intron (not found in other eukaryotes including other Excavata species), and thus the four introns all are ‘relatively new’ introns in ‘relatively new’ genes (Table 1).

**Table 1.** The integrated information of the eight spliceosome-mediated cis-spliced introns and their host genes and complementary strands.

| Intron-containing gene |  |  | Intron |  | Complementary strand |
| --- | --- | --- | --- | --- | --- |
| Gene ID | Product | Age | Age | Size (bp) |  |
| GL50803-27266 | 2Fe-2S ferredoxin | ancient | relatively new | 35 | No ORF, no transcripts detected |
| GL50803-15604 | 26S proteasome non-ATPase regulatory subunit 4 |  | ancient | 29 | No ORF, no transcripts detected |
| GL50803-15124 | Dynein light chain |  |  | 32 | No ORF, no transcripts detected |
| GL50803-17244 | Ribosomal Protein L7A |  |  | 109 | **Antisense gene with transcripts** |
| GL50803-37070 | Hypothetical protein | relatively new | relatively new | 41 | **Antisense gene with transcripts** |
| GL50803-35332 | Hypothetical protein |  |  | 220 | No ORF, no transcripts detected |
| GL50803-15525 | Hypothetical protein |  |  | 33 | No ORF, no transcripts detected |
| GL50803-86945 | Hypothetical protein |  |  | 36 | No ORF, no transcripts detected |

These observations suggest that: 1) while *Giardia* massively lost its introns, new introns also arose in both the ancient and relatively newly evolved genes; 2) the pressure of intron loss seems to constantly exist in the whole evolutionary process of *Giardia*, but only a few of both the ancient and newly-emerged introns have been retained in the genome.

To find the reason why these few introns can be retained in *Giardia* genome under strong pressure of loss, we investigated the characteristics of these introns in many aspects. It had been shown that seven of the eight *Giardia* introns are bounded by canonical GT-AG splice signals, only one, the [2Fe-2S] ferredoxin (GL50803-27266) intron, has a noncanonical splice signal CT-AG [19]. The sizes of the eight introns are all small (most of them are less than 40 bp long and are not the multiple of three) and do not have any conserved sequence motifs. Our further analysis predicted no special secondary structures that would be able to form in these introns. Besides, our survey also showed that there were not any reports about alternative splicing of the two introns in genes GL50803-15525 and GL50803-86945 [18, 22]. Taken together, these results suggest that the retention of the few introns seems to be neither due to the structural features nor necessarily due to the functional importance of these introns.

Interestingly, on the complementary strands, we found that two intron-containing genes, GL50803-17244 (ribosomal protein L7a gene) and GL50803-37070 (a “hypothetical protein” gene), each have an antisense gene, GL50803-20429 and GL50803-28204, which are just annotated as “hypothetical protein” and “unspecified product” in the genome database, respectively. That is, the two intron-containing genes and their antisense gene form SAS gene pairs. We thought this phenomenon might be related to the intron retention. Nevertheless, the two anti-sense genes need to be further verified, and the details of the overlaps with their sense genes also need to be analyzed in detail.

### Verification of the antisense genes

The strand-specific RT-PCR of the two antisense genes, GL50803-28204 and GL50803-20429, generated two products with the expected lengths of 172 and 288 bp, respectively. The sequencing further confirmed that the two products just seem to be transcribed from the opposite direction of the two sense (intron-containing) genes, GL50803-37070 and GL50803-17244, respectively, and the two introns are just located within the two overlapping regions of the two SAS gene pairs, respectively (Figure 1).

**Figure 1.**
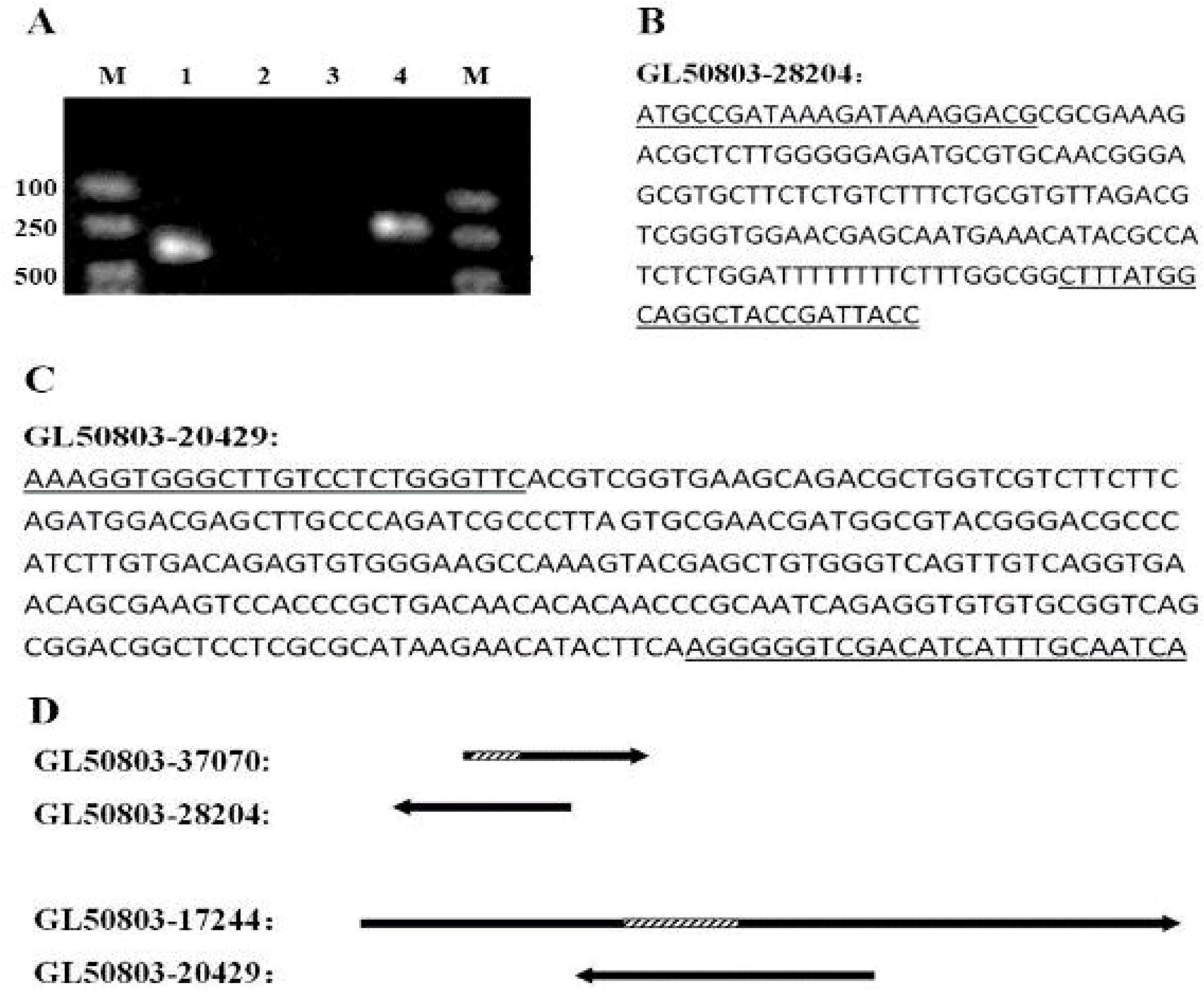
Results of strand-specific RT-PCR and sequencing of the two antisense genes, and the schematic diagram of this two SAS gene pairs. **A.** Lane 1, Strand-specific RT-PCR product of the GL50803-20429; Lane 4, the Strand-specific RT-PCR product of the GL50803-28204; Lane 2 and lane 3, negative controls (with no RTase) corresponding to lane 1 and lane 4, respectively; M, molecular markers. **B.** Nucleotide sequence of GL50803-28204 gene acquired by Strand-specific RT-PCR and sequencing. The locations of the primers are underlined. **C.** Nucleotide sequence of GL50803-20429 gene acquired by Strand-specific RT-PCR and sequencing. **D.** Schematic diagram of the two SAS gene pairs. The sequence lengths of GL50803-28204 and GL50803-20429 are according to the Strand-specific RT-PCR products, and the lengths of GL50803-37070 and GL50803-17244 are based on the GiardiaDB database. Arrow represents the orientation of transcription; and the dashed box and solid lines represent introns and exons, respectively.

The RACE of the two antisense genes generated a 1232 bp product for GL50803-28204 and a 1177 bp product for GL50803-20429. After sequencing and comparing with their corresponding genomic DNA sequences in the GiardiaDB database, we found the two antisense genes contain no introns, especially in the regions corresponding to the two introns of the sense genes. But the software ORF Finder predicted that the largest ORFs of the two antisense genes are only 264 bp for GL50803-28204 and 363 bp for GL50803-20429, and moreover, no proteins homologous to the putative proteins coded by the two largest ORFs could be found in other organisms including *Giardia’s* close relative *Spironucleus* in GenBank. Therefore, the two antisense genes have transcriptional activity and are most likely *Giaridia-specific* non-coding genes.

## Discussion

To investigate the reasons for the evolutionary retention of the few introns in the eukaryotes having undergone massive intron loss, we chose the extremely intron-poor eukaryote *Giardia* as the study object. When investigating the characteristics (including distribution patterns) of the eight intron-containing genes and their corresponding introns in *Giardia* genome, we found that in spite of the massive loss of introns, intron gain also occurred in both *Giardia’s* ancient and relatively newly-evolved genes; it turns out that the selective pressure of intron loss seems to constantly exist in the whole evolutionary course of *Giardia*, but a few of both the ancient and the newly-emerged introns have been retained in modern *Giardia*.

To explore how these few introns can be retained under the constant strong pressure of loss. First, we investigated whether there exist some features in these few intron-containing genes and the introns probably responsible for the retention, but failed to find any special regularities or unusualnesses in many aspects such as splicing signal, intron size and secondary structure, and alterative splicing. This suggests that there are neither any special structural features nor necessarily any functional importance of introns responsible for the intron retention. This is consistent with the fact that so many introns, at least part of which definitely possesses important functions, have been lost in intron-poor eukaryotes like *Giardia*. Thus, the reasons for the retention might lie outside the intron-containing genes and the introns themselves.

Interestingly, we noticed that on the complementary strands of two of the eight intron-containing genes, GL50803-37070 and GL50803-17244, there exist correspondingly two anti-sense genes, GL50803-28204 and GL50803-20429, though they are just annotated as “product unknown” in GiardiaDB. By strand-specific RT-PCR, RACE and sequencing, we got the transcripts and sequences of the two genes and found they both have no introns. Thus the two anti-sense genes have been verified to be really genes that are actively transcribed. And actually the anti-sense gene GL50803-20429 has been reported to be a mRNA gene being expressed during excystation and encystation, and in trophozoites but not cysts [23]. As for the other anti-sense gene GL50803-28204, it has a quite short putative ORF but has no homologs in other organisms including *Spironucleus*. Although the corresponding DNA sequence regions in the four other *G. lamblia* isolates (DH, P15, GS and GS-B) with genomic data exhibit significant similarities (≥ 83.9% similarity) to those of the two anti-sense genes, there are not any annotations and transcriptome information about those regions. Besides, the total RNA were processed using Poly(A) Polymerase to add a poly(A) tail at the 3’ends before we performed rapid amplification of their cDNA 3’ends, thus from the experiment we still did not know whether the transcripts of GL50803-28204 are polyadenylated or not, namely, mRNA or not. Therefore, we can only conjecture that the two anti-sense genes are either non-protein-coding genes or Giardia-specific protein-coding genes. Considering that many identified non-coding RNAs in *Giardia* overlap with protein-coding genes on the antisense strands [24], the antisense gene might also be noncoding RNA gene. But there is still no tangible evidence for what the two genes are despite our lots of experimental efforts (not shown) to identify them. Whatever the antisense genes code for, our work showed that they are functional genes and form SAS gene overlap with their intron-containing sense genes, and that the introns just reside in the overlap regions. Considering that the gene sequence mutation (especially deletion) cannot occur randomly, the antisense genes must have imposed the restriction of variation (especially deletion) on the introns of the sense genes in the overlapping areas, and thus such a kind of SAS gene overlap must have prevented the introns from being lost.

As for the other six introns, we did not find any ORFs on their corresponding complementary strands. Although we also experimentally examined whether their complementary strands (especially the areas overlapping with these introns) are transcribed, no transcripts were found (Additional file 2). Nevertheless, it is uncertain whether the corresponding complementary strands of these introns are resided by some unknown genomic functional elements which are not transcribed at all. If this is true, these introns are also retained by the same cause as the former two ones. Certainly, it is also possible that the six introns are retained by other unknown reasons. We have also analyzed the intron regions of many intron-poor eukaryotes including *Microsporidia, Trichomonas, Spironucleus*, but unfortunately did not find such sense-antisense gene overlaps as in *Giardia* (data not shown). But considering some genes overlap with the UTRs of the adjacent genes (as found in *Antonospora locustae* and *Encephalitozoon cuniculi* [25]), we can not obtain the UTRs information of those genes in current database, thus many of the overlaps might be able to be found out. More importantly, some introns might overlap with unknown genomic functional elements including non-coding and non-transcribed ones, since there are so huge remaining component of eukaryotic genomes, much of which was traditionally regarded as “junk” and is still undetermined. This might be the important cause for that few SAS gene overlaps in intron regions that can be identified at present.

Theoretically, overlapping with any genomic functional elements on either the same strand or the complementary one (namely, either same-strand overlapping or different-strand overlapping) can result in intron retention, as long as the introns are just in the overlapping areas. Therefore, since such overlapping structures are widely distributed in eukaryotes [26], it can be expected that quite a number of introns in diverse eukaryotes may also be retained due to this kind of “overlap constraint”. We believe that more and more examples might be able to be found in diverse eukaryotes in the future. Conversely, such an intron retention phenomenon probably can provide a valuable clue to find new genomic functional elements – in the overlapping area with introns.

## Conclusions

In conclusion, by investigating the extremely intron-poor eukaryote *Giardia*, we have revealed some interesting findings about the dynamic evolution of introns in the intron-poor eukaryotes: the pressure of intron loss may constantly exist in these eukaryotes, but new introns can still arise either in ancient genes or new-evolved genes, but only a few introns can be retained in the genome; the retention of the few introns is not caused by special features or functional constraint of the introns themselves but due to the reasons outside of the introns, and “overlap constraint” imposed by other genomic functional elements overlapping with the introns is at least an important one of the causes. First, our findings not only support the “intron-rich ancestor” theory, but also can explain why few eukaryotes were found to be completely intronless. Second, our finding may conversely provide a clue to find new genomic functional elements (which was probably traditionally regarded as “junk” and is still undetermined) in such kinds of overlap regions. Most importantly, our work implicates that “functional constraint” of introns is not necessarily directly associated with intron loss and gain, or that the real functions or the way of functioning of introns are probably still outside of our current knowledge. Therefore our work may be able to shed some new lights on the research of evolution and function of introns and genomes.

## Methods

### Database and bioinformatics methods

The template sequences for designing primers of *Giardia* genes were downloaded from GiardiaDB (December 1, 2017 release) [27]. The software ORF Finder were used to predict the ORFs of the RACE products (see blow), then the predicted ORFs were used as queries to search their homologs with Blastp against the NCBI non-redundant protein sequences (nr) database. The program RNAfold web server was used to analyze the secondary structure of introns. The sequences of the four genes and their coding proteins, 2Fe-2S ferredoxin, 26S proteasome non-ATPase regulatory subunit 4, Dynein light chain, and Risbosomal Protein L7A from other organisms were identified and collected by Blastp searching against GenBank with *Giardia’s* corresponding sequences as queries. Protein alignments were generated with ClustalX 2 applying default alignment parameters. The introns in the genes were determined by comparing cDNA and gene sequences. The other four intron-containing genes with annotated as hypothetical protein also identified by sequence comparative analysis to determine whether they are Giardia-specific or not.

### *Giardia* cultures

The cell line of WB isolate (assemblage A), namely WB clone C6 (ATCC 50803), was used in the study. Its cultures were routinely grown in filter-sterilized TYI-S-33 medium supplemented with bovine bile in glass screw cap tubes at 37 °C and were sub-cultured every 3 to 4 days.

### RNA extraction

*Giardia* total RNA was extracted and treated to remove any contaminated genomic DNA by RNAprep Pure Cell/Bacteria Kit (TIANGEN) using about 5×10^6^ *Giardia* trophozoites that were harvested by ice-slush incubation and centrifuged at 6000g for 5 min according to the manufacturer’s instructions. The quality and quantity of the RNA preparation were assessed with agarose gel electrophoresis and the absorption at 260 and 280nm.

### Strand-specific RT-PCR

First-strand cDNAs of the two antisense genes, GL50803-28204 and GL50803-20429, were synthesized from 500ng DNase-I treated total RNA per reaction at 54°C 30 min, 99°C 5°C min, and 5°C 5 min with the specific antisense primer 4C and 9C respectively instead of the reverse primers in the kit, and then amplified with specific primers pairs of 4A/4S and 9A/9S by using RNA PCR Kit (AMV) Ver.3.0 (Takara, Japan). The PCR conditions were as follows: 94°C for 30 s, followed by 30 cycles of 94°C for 30 s, 55°C for 30 s and 72°C for 45s. The PCR products were purified using the Wizard SV Gel and PCR Clean-Up System kit (Qiagen, Germany), and cloned into pMD-19T vectors from 35 ng purified PCR products using TaKaRa pMD-19T Vector Cloning Kit (TaKaRa, Japan) according to the manufacturer’s instructions. Then, the ligation products were transformed into DH5 Chemically Competent *E. coli*. Colony PCR with vector-specific primers provided in the kit was adopted to select colonies. These selected colonies were sequenced using vector-specific forward and reverse primers by Sangon Biology Company (Shanghai, China).

### Rapid amplification of cDNA ends

The total RNA were processed using Poly(A) Polymerase(TaKaRa, Japan) to add a poly(A) tail at the 3 ends of the RNA before performing rapid amplification of their cDNA 3’ends. We experimentally determined the 3’ends by using nested PCR primer (3R4O/3R4I and 3R9O/3R9I) according to the RNA PCR Kit (AMV) Ver.3.0 (Takara, Japan). 5’-RACE was performed by using a SMARTer RACE 5’/3’Kit (TaKaRa, Japan) with 500ng total RNA as the template and the gene-specific 5’-RACE primers 5R4 and 5R9 for the two antisense genes, GL50803-28204 and GL50803-20429, according to the manufacturer’s instructions. Both the 3’-RACE and 5’-RACE primers (Additional file 3) were designed based on the transcripts from the Strand-specific RT-PCR. The RACE-PCR products were analyzed by agarose gel electrophoresis and sequenced as described above.

## List of abbreviations

LECA: : last eukaryotic common ancestor.
SAS: :sense-antisense.
RACE: : rapid amplification of cDNA ends.

## Declarations

### Ethics approval and consent to participate

Not applicable

### Consent for publication

Not applicable

### Availability of data and materials

All data generated or analysed during this study are included in this published article and its supplementary information files.

### Authors’ contributions

J.W. designed and supervised this study. M.X performed genetic characterization work. M.X., B.C., Q.Y., J.S., and Z.L. analyzed the data. M.X. and J.W. wrote the manuscript. All authors read and approved the final manuscript.

## Acknowledgements

Not applicable

## Additional files

**Additional file 1:** The conservative analysis of Giardia’s introns among diverse eukaryotes

**Additional file 2:** Results of strand-specific RT-PCR of the complementary areas of the other six introns of *G. lamblia*.

**Additional file 3:** The primers designed for the strand-specific RT-PCR and RACE of the complementary areas of the eight introns of *G. lamblia*.

